# Robustness and adaptability of sensorimotor skills in expert piano performance

**DOI:** 10.1101/2023.10.02.560469

**Authors:** Masaki Yasuhara, Kazumasa Uehara, Takanori Oku, Sachiko Shiotani, Isao Nambu, Shinichi Furuya

**Affiliations:** Nagaoka University of Technology, Nagaoka, Japan; Sony Computer Science Laboratories Inc., Tokyo, Japan; Department of Computer Science and Engineering, Toyohashi University of Technology, Toyohashi, Japan; NeuroPiano Institute, Kyoto, Japan

**Keywords:** feedback control, plasticity, sensorimotor integration, music

## Abstract

Skillful execution of sequential actions requires the delicate balance of sensorimotor control, encompassing both robustness and adaptability. Previous studies have characterized behavioral and electrophysiological responses to sensory perturbation during performance of sequential movements such as speech and singing. However, it remains unknown whether and in what manner both motor and neural responses, triggered by sensory perturbation, undergo plastic adaptation as a consequence of extensive sensorimotor experience. Here, we addressed this question by comparing effects of transiently delayed tone production on the spatiotemporal patterns of the subsequent motor actions and event-related potentials (ERPs) during fast and accurate piano performance between expert pianists and musically-untrained individuals (non-musicians). Following the delayed tone production, the inter-keystroke interval was abnormally prolonged in non-musicians but not in pianists. By contrast, the keystroke velocity following the tone delay was increased only in the pianists. A regression model further demonstrated that the change in the inter-keystroke interval following the perturbation covaried with the ERPs of the N180 and P300 components particularly at the frontal and parietal regions. In contrast, the alteration in the keystroke velocity was associated with the P300 component of the temporal region ipsilateral to the moving hand, which suggests enhancement of auditory but not somatosensory feedback gain following auditory perturbation. Together, these findings suggest that distinct neural mechanisms underlie robust and adaptive sensorimotor skills individuals with different levels of proficiency.

## Introduction

Skillful behaviors are typically characterized by harmonizing both robustness and adaptability of sensorimotor control. A challenge in fast and accurate performance of sequential motor actions such as speech, typing, and musical performance is to accommodate uncertainty originating from the stochastic biological system (e.g. sensorimotor noises) (Osborne et al., 2005; Faisal et al., 2008) and unpredictable perturbation from the environment (Leonardo and Konishi, 1999; Tremblay et al., 2003; Sober and Brainard, 2009). The nervous system is therefore required to optimally integrate predictive and adaptive control of movements so as to fulfill task requirements under uncertainty (Diedrichsen et al., 2010; Scott, 2016; Uehara et al., 2023). One approach to probe into this mechanism is to provide artificial sensory perturbation during motor actions (Howell and Archer, 1984; Desmurget et al., 1999; Sainburg et al., 1999; Furuya and Soechting, 2010; Tian and Poeppel, 2015; Popp et al., 2022). For instance, continuously delaying the timing of tone production in speech and musical performance commonly disrupts the ongoing motor actions (Pfordresher and Palmer, 2002; Hashimoto and Sakai, 2003; Ozker et al., 2022). However, most of previous studies have focused on motor reaction to sensory perturbation only in well-trained tasks, which limits the understanding of neuroplastic mechanisms subserving skillful sensorimotor control responsible for behavioral stability and adaptability.

Musical performance can be suitable for addressing this issue. To compare effects of sensory perturbation on motor actions between musicians and musically-untrained individuals (i.e. non-musicians) have unveiled specialized sensorimotor skills in relation to expertise (Zatorre et al., 2007; Hirano et al., 2020). Neural and behavioral responses to altered auditory or somatosensory feedback in musical performance differed between musicians and non-musicians (Jones and Keough, 2008; Zarate and Zatorre, 2008; Kleber et al., 2013; van der Steen et al., 2014). For example, following a transient delay of timing of tone production in piano playing, non-musicians but not expert pianists, abnormally slowed down the local tempo, exhibiting movement disruption (van der Steen et al., 2014). In addition, the amount of the movement disruption to the perturbation was positively correlated with the age at which pianists commenced their musical training. In contrast, pianists but not non-musicians struck the key harder in response to the perturbation, which allows for elevating either somatosensory or auditory gain in motion. Yet, it has not been known what neural mechanisms mediate expertise-dependence of robust and adaptive control of fast and accurate production of sequential motor actions. One candidate neural signature is the event-related potentials (ERPs) that emerge in response to transient sensory perturbation (Maidhof et al., 2010; Strübing et al., 2012; Sammler et al., 2013; Bianco et al., 2016). Transient alteration of the pitch in sequential production of piano tones evoked error-related negativity at the fronto-central region, which was more pronounced during playing the piano than listening (Maidhof et al., 2010). Based on the behavioral observation, it is plausible that the ERPs evoked during piano performance depend on musical proficiency. For instance, a cortical region eliciting ERPs that is associated with adaptive motor response to delayed tone production lets us identify whether movement flexibility in response to sensory perturbation is associated with gain modulation of unperturbed or perturbed sensory modality. A bottleneck to test it is no established clue to assess neurophysiological responses to sensory perturbation during fast skillful behaviors.

Here we addressed this issue by developing a novel system that simultaneously provides sensory perturbation and assesses behavioral and neurophysiological responses during fast piano performance in high temporal resolution. Our high-speed sensing system that measures piano key motions (Oku and Furuya, 2022) in synchronization with the measurement of electroencephalogram (EEG) was capable of comparing effects of transiently delayed production of a piano tone on the sequential finger movements and electrophysiological activities between expert pianists and non-musicians. By leveraging it, we characterized expertise-dependent behavioral and electrophysiological responses to sensory perturbation as well as their relationship during piano performance.

## Materials and Methods

### Participants

Fifteen expert pianists (mean ± standard deviation of age was 25.5 ± 7.2 years, 10 females) and 15 non-musicians (mean ± standard deviation of age was 25.6 ± 4.8 years, 10 females) participated in the study. All pianists started to play the piano at 4.6 ± 1.2 years old (mean ± SD across participants), and had specialized piano education at music conservatories, whereas the non-musicians had no piano training or less than three years of piano training. The local ethics committee of Sony Corporate approved this study in accordance with the guidelines established in the Declaration of Helsinki. Before starting the data collection, we obtained written informed consent from all participants.

### Experimental setup

A custom-made experimental system was developed in order to synchronously record EEG data and data on piano keystrokes (Fig. 1A). A digital piano (VPC1; KAWAI, Hamamatsu, Japan) that implemented position sensors under all piano keys (so called “HackKey”) (Oku and Furuya 2022) enabled to measure the time-varying vertical position of piano keys. The core experimental program was operated by Psychopy (Peirce et al. 2019) (Fig. 1A). The program provided the instructions to participants using a display connected to the PC. The HackKey thread was responsible for monitoring and recording key vertical position over time at a sampling rate of 1kHz, and once detecting a key press that amounts 5 mm, a trigger signal was sent to the EEG recording device via an NI-DAQ (National Instruments Corporation, Texas, US). In parallel, the time-varying key position data were stored for each session. The MIDI thread either passed through the input MIDI signals or artificially delayed the timing of issuing the output signal by 120 ms (i.e. delayed auditory feedback). A Rubix22 (Roland, Hamamatsu, Japan) was used for inputting and outputting the MIDI signals. The MIDI signals processed by the MIDI thread were converted to piano sounds by a Sound Processor (PIANO BOX PRO; MIDITECH, Cologne, Germany). Participants were provided with auditory feedback through earphones (MDREX155; SONY, Tokyo, Japan) connected to the PIANO BOX PRO.

**Fig. 1.**
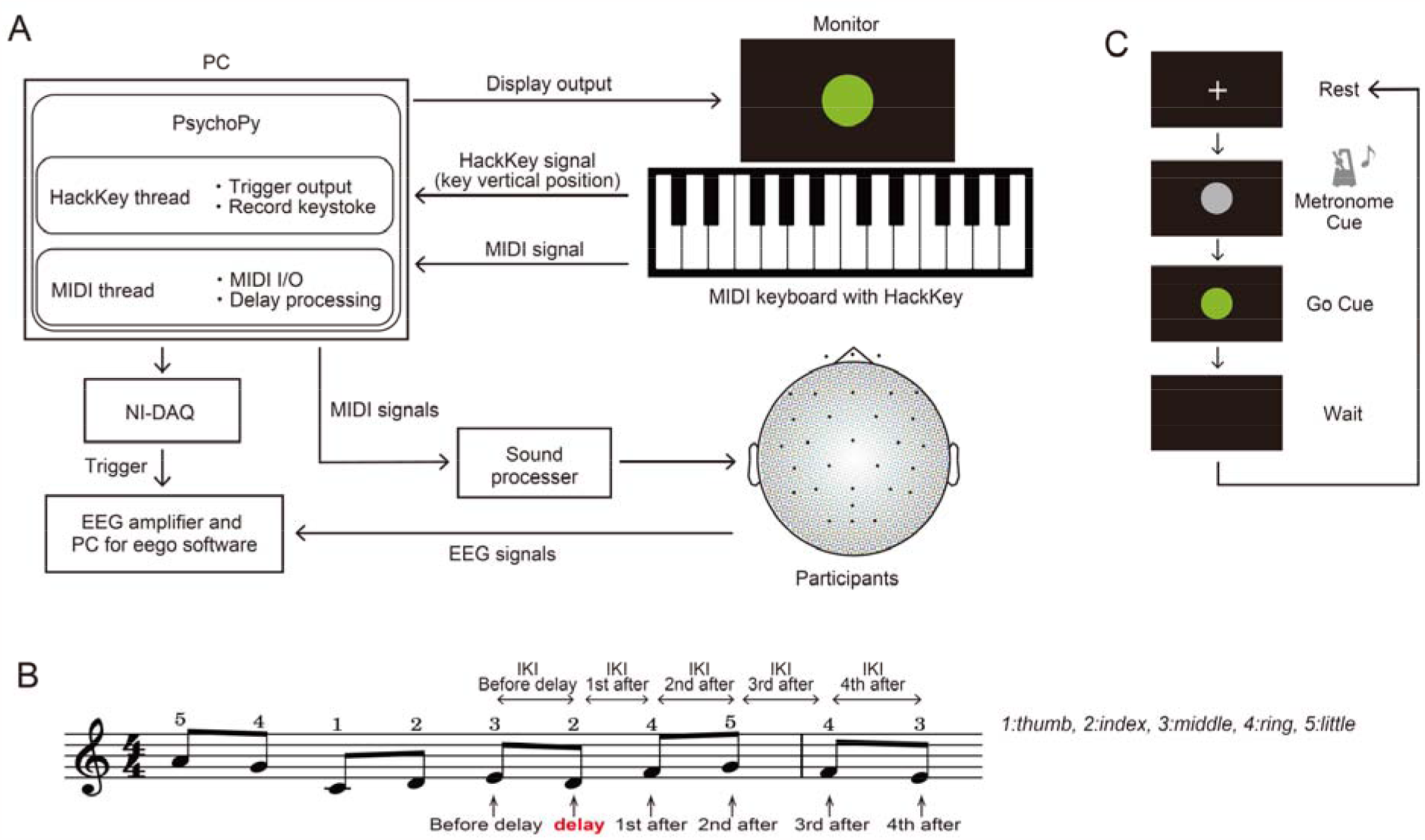
Experimental setup. (A) The architecture of the system for data recording and stimulus presentation. When a participant pressed the keys on the piano keyboard, the MIDI signal was transmitted into the computer (PC), and the generated sound was fed back into the participant through the binaural earphones. In parallel, the time-varying data of the key vertical position was measured by 1kHz with the HackKey system implemented in the keyboard as “HackKey signal”. When the key position first moved over 5 mm, the trigger signal was sent to the EEG amplifier for the synchronous recording. The EEG signals were recorded at 1kHz from a 32-channel cap throughout a session. (B) Musical score was presented to the participants prior to performing the experiment, which designated the sequence of tones and fingering. Information on the amount of timing delay of tone production was not provided. Although the musical score was not displayed on the monitor, the participants were allowed to read the score whenever necessary. The second D4 note (i.e. 6^th^ tone) involved delayed tone production by 120 ms with a 30% probability. IKI: the inter-keystroke interval, VEL: the key descending velocity. (C) An experimental pipeline. Participants played the piano following the instructions displayed on the monitor. Before the Go cue was provided, a metronome sound at either 80 BPM or 160 BPM was randomly presented, and participants were instructed to play at the designated tempo.

### Experimental design

Participants were asked to play a sequence of ten tones requiring the use of all five digits of the right hand (Fig. 1B). The system delayed the timing of tone production by 120 ms with a 30% probability on the second strike of the D4 key. Prior to the performance, a visual cue was presented to the participants via a display placed in front of them, and a metronome provided tones prior to the GO cue, indicating the tempo to be played with (Fig. 1C). Metronome tones at two tempi were randomly provided, either 80 beats per minute (BPM) or 160 BPM, and participants were instructed to perform according to the played tempo when the Go cue was presented. The target inter-keystroke interval was 375 ms for 80 BPM (= intermediate tempo) and 187.5 ms for 160 BPM (=fast tempo), respectively. The experiment consisted of two sessions, each with 100 trials, which included 30 trials with the provision of delayed auditory feedback (DAF) for each of the two tempi. The participants were instructed to play as accurately as possible at the target tempo with legato touch. The participants were also asked to keep watching the display put in front of them without seeing their hands and piano keys.

### EEG recording

Continuous EEG data was recorded using EEG equipment (eego sports, ANT Neuro, Enschede, The Netherlands) with a 32-channel electrode cap (WaveGuard EEG cap, Advanced Neuro Technology, Netherlands) in accordance with the international 10-10 layout at a sampling rate of 1kHz. The ground and system reference electrodes were placed at AFz and CPz, respectively. Skin/electrode impedance was kept at below 5 kΩ throughout data collection. The EEG signal was interpolated with the surrounding electrodes throughout off-line analysis using the “evoked.interpolate_bads” function in the MNE-Python when a high impedance electrode was detected.

### Data analysis

#### Behavioral responses

During the session, the time-varying key position data were recorded. Using this data, the inter-keystroke interval (from one key depression to the subsequent key depression) and the peak key descending velocity were computed. The key depression was defined as an event when a key moved down by 5 mm from the neutral position. The changes in the keystroke timing and loudness in response to the transient tone delay were defined as the difference in the average values of each measure between the perturbed and unperturbed conditions. The trials of the unperturbed condition were therefore used as a baseline.

#### Event-related potentials

MNE-python (Gramfort et al., 2013) was used for the EEG data analysis. The EEG data were first re-referenced to the common averaged reference. A high-pass filter with a cutoff frequency of 1 Hz was applied to remove linear trends and a notch filter targeting 50 Hz (49.75–50.25 Hz) was used to eliminate power line noise. Artifacts arising from eye blinks and muscle contractions were excluded using independent component analysis (ICA) with the InfoMax algorithm. The excluded components were selected based on ICLabel (Pion-Tonachini et al., 2019) which was implemented in Python-based software (Li et al., 2022). ICA-processed EEG data were epoched ranging from -100 to 600 ms with respect to the trigger onset (defined as time 0). Epochs including residual artifacts, were detected using an EEG amplitude criterion (above 100 μV) and were then excluded from the reported results. In addition, the erroneous performances were also detected and excluded according to the recorded trigger signal produced upon each key press. Grand averages were then calculated for each condition (presence or absence of delayed events, tempi), and event-related potentials (ERPs) were obtained from each individual. The changes in grand-averaged ERPs in response to the tone delay were defined as the difference in the average values of each measure between the perturbed and unperturbed conditions. The trials in the unperturbed condition were treated as a baseline. To quantify the amplitudes and latencies of ERPs, we selected a first component of 80 ms in length (140– 220 ms) for N180, and a second component of 140 ms in length (230–370 ms) for P300. These windows were selected based on our observation of ERPs (see Fig. 3) and the previous study (Knolle et al., 2013) demonstrated that the observations of N180 and P300 represent the neurophysiological response to auditory stimuli.

### Statistics

The whole statistical analyses for the behavioral and EEG responses were performed using R (version 4.4.3) and glmnet package (version 4.1.7). Post hoc tests with the Benjamini-Hochberg correction for multiple comparisons were performed in the case of significance. A statistical significance level was defined as p = 0.05.

#### Behavioral responses

To test the effects of the perturbation on the subsequent movements, a three-way mixed-design analysis of variance (ANOVA) with a group (pianists and non-musicians) as a between-subject variable, and tempo (80 BPM and 160 BPM), and event (strikes or intervals before, during and after the delayed tone production, see Fig. 2) as within-subject variables were carried out. If the assumption of sphericity was violated, the Greenhouse-Geisser correction was performed. Post-hoc tests were performed to test differences between groups and tempi. In addition, two-tailed t-tests were performed to test a difference from zero (=baseline).

**Fig. 2.**
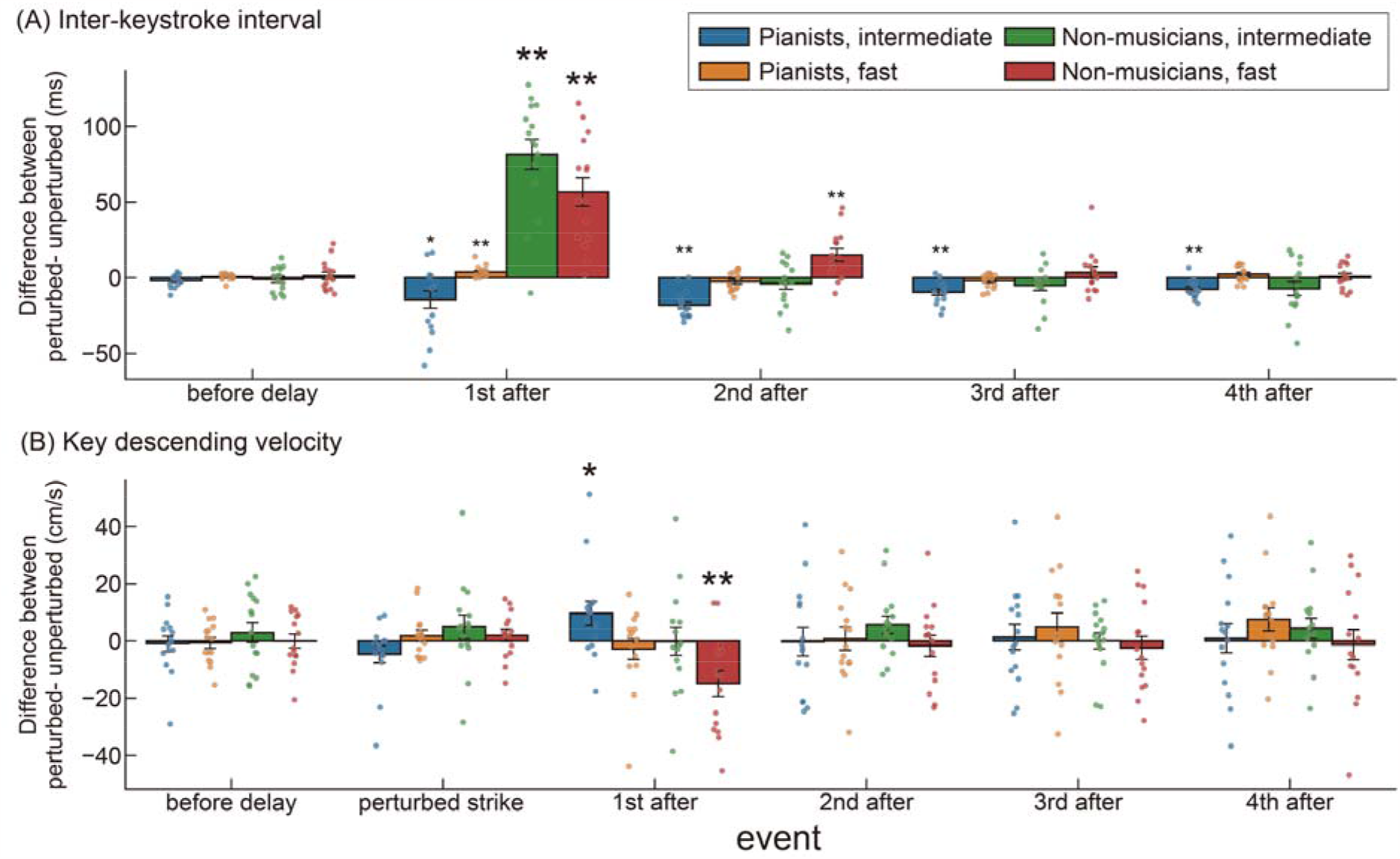
Effects of the transient delay of tone production on keystroke during piano playing. An error bar indicates one standard error across the participants. Blue, orange, green, and red boxes represent expert pianists at the intermediate and fast tempo, non-musicians at the intermediate and fast tempo, respectively. The value represents a difference between the conditions with and without the tone delay at each event. (A) inter-keystroke interval and (B) peak key descending velocity. The asterisk indicates the value with a significant difference from zero. Statistical significance: *p < 0.05, ** p < 0.01. See Fig. 1B for the description of the events.

#### Event-related potentials

To test the effects of delayed auditory feedback on ERPs, a three-way mixed-design ANOVA was performed with group (expert pianists and non-musicians), tempo (80 BPM and 160 BPM), and EEG channel (six representative channels; Fz, Cz, Pz, Oz, T7, T8) as independent variables. To avoid massive multiple comparison problems associated with post-hoc testing after ANOVA (Groppe et al., 2011), we opted for 6 representative EEG channels of each brain region (frontal, bilateral temporal, central, parietal, occipital) from 32 channels. If the assumption of sphericity was violated, the Greenhouse-Geisser correction was applied. Post-hoc tests were performed to further test differences between groups and tempo factors.

#### Multiple regression analysis

The primary goal of this study was to identify neural correlates of the robustness and adaptability to the transient auditory perturbation. To infer how brain activities are associated with behavioral responses, we performed a penalized multiple regression analysis with L2 norm regularization, so called Ridge regression, including all pianists and non-musicians in the model. Ridge regression was used to avoid the problem of multicollinearity. The λ parameter that determines the overall intensity of regularization was optimized through the leave-one-subject-out cross-validation to resolve over-fitting problems. The goodness of the fitting model was expressed by the R squared value (R^2^). Based on the findings from our previous study (van der Steen et al., 2014) and from the results of behavioral responses (see Fig. 2), we used three behavioral responses as dependent variables, which are the first and second inter-keystroke intervals after the perturbation and the first key descending velocity after the perturbation. The inter-keystroke interval following the delayed tone production indicates whether or not the performance temporarily slowed down, representing the robustness of tempo control to the perturbation. The first key descending velocity after the perturbation indicates whether the keystroke becomes stronger, which is considered to be a reaction to adapt to the perturbation (Furuya and Soechting, 2010; van der Steen et al., 2014). In this regression analysis, we focused on the intermediate tempo condition data, because both the shorter interval and stronger keystroke following the delay in the pianists were evident only at this tempo condition.

## Results

We excluded 31.4% of the trials from the analyses across all participants due to excessive noise in the preprocessed EEG signals and/or erroneous performance.

### Behavioral responses

The mean changes in both the inter-keystroke interval and the peak key descending velocity at different events related to the delay-manipulation were computed to assess the effect of delayed auditory feedback on accuracy of the subsequent movement production (Fig. 2). The value of the keystroke prior to the delay functions as a baseline of the normal unperturbed movement production.

ANOVA showed a significant three-way interaction effect for the mean inter-keystroke interval response (Table 1). Post hoc tests revealed the first inter-keystroke interval after the delayed tone production differed between the pianists and non-musicians at both the intermediate tempo (F (1,28)=70.2, p<0.05) and fast tempo (F(1,28)=32.1, p<0.05). Similarly, the second inter-keystroke interval differed between the groups at both the intermediate tempo (F (1,28) =10.6, p<0.05) and fast tempo (F(1,28)=15.7, p<0.05). For both the first and second inter-keystroke intervals after the perturbation, the difference between the perturbed and unperturbed conditions was smaller for the pianists compared to non-musicians. Post hoc test also revealed that the inter-keystroke interval in the pianists differed between the two tempi at the first four successive strikes following the perturbation and that the difference between the perturbed and unperturbed conditions was smaller at slower tempo. For the non-musicians, the inter-keystroke interval differed at the first and second strikes following the perturbation between the two tempi. The one-tailed t-tests revealed that the response of the inter-keystroke interval at the first strike after the perturbation was significantly greater than zero in the non-musicians (p<0.05), confirming the prolongation of the local tempo in response to the disruption. For the pianists, by contrast, the response of the inter-keystroke intervals to the perturbation was significantly smaller than zero at the intermediate tempo, confirming continuous speed-up of the local tempo.

**Table 1.** Results of the three-way mixed-design ANOVA for the behavioral responses (i.e. local tempo and keystroke velocity) to the pertubation.

|  |  | Interval | Velocity |
| --- | --- | --- | --- |
| Group | F(1,28) | 34.920 | 0.488 |
|  | p | **2.34x 10<sup>-6</sup>** | 0.491 |
|  | eta<sup>2</sup> | 0.284 | 0.003 |
| Tempo | F(1,28) | 12.545 | 0.681 |
|  | p | **0.001** | 0.416 |
|  | eta<sup>2</sup> | 0.046 | 0.008 |
| Event | F(4,112) | 57.531 | 0.925 |
|  | p | **1.61x 10<sup>-10</sup>** | 0.448 |
|  | eta<sup>2</sup> | 0.449 | 0.01 |
| Group x Tempo | F(1,28) | 4.672 | 1.248 |
|  | p | **0.039** | 0.273 |
|  | eta<sup>2</sup> | 0.018 | 0.014 |
| Group x Event | F(4,112) | 61.183 | 2.916 |
|  | p | **7.19x 10<sup>-11</sup>** | **0.027** |
|  | eta<sup>2</sup> | 0.465 | 0.032 |
| Tempo x Event | F(4,112) | 8.206 | 4.616 |
|  | p | **0.000228** | **0.004** |
|  | eta<sup>2</sup> | 0.049 | 0.031 |
| Group x Tempo x Event | F(4,112) | 13.388 | 0.539 |
|  | p | **2.23x 10<sup>-6</sup>** | 0.665 |
|  | eta<sup>2</sup> | 0.077 | 0.004 |

Interval: inter-keystroke interval
Velocity: key descending velocity
A bold number indicates a significant effect (p<0.05)

For the mean change in the key descending velocity, there were significant interaction effects between group and event, and between tempo and event (Table 1). The post hoc tests revealed the first key velocity after the tone delay significantly differed between the pianists and non-musicians and that the difference between the perturbed and unperturbed conditions was larger for the pianists. For the tempo, post hoc tests revealed the first key velocity after the perturbation differed between the two tempi. The one-tailed t-test revealed that the change in the key descending velocity was significantly smaller than zero for the non-musicians when playing at fast tempo, indicating the softer keystroke. Conversely, in pianists, the keystroke velocity after the perturbation was larger than zero when playing at the intermediate tempo, indicating harder keystroke following the tone delay.

Overall, these behavioral findings indicate that the piano performance was influenced by transiently delayed auditory feedback in a distinct manner between the pianists and non-musicians. The differences between the pianists and non-musicians were evident, displaying smaller rhythmic disruption of motions and stronger strike following the perturbation in the more skilled group.

### ERPs responses

To investigate ERPs elicited by the delayed tone production, the changes in grand-averaged ERPs were computed by subtracting the value at the unperturbed condition from one at the perturbed condition (Fig. 3). The negative component N180 was elicited at around 180 ms following the keystroke, and the positive component P300 was elicited at around 300 ms in both of the groups.

**Fig. 3.**
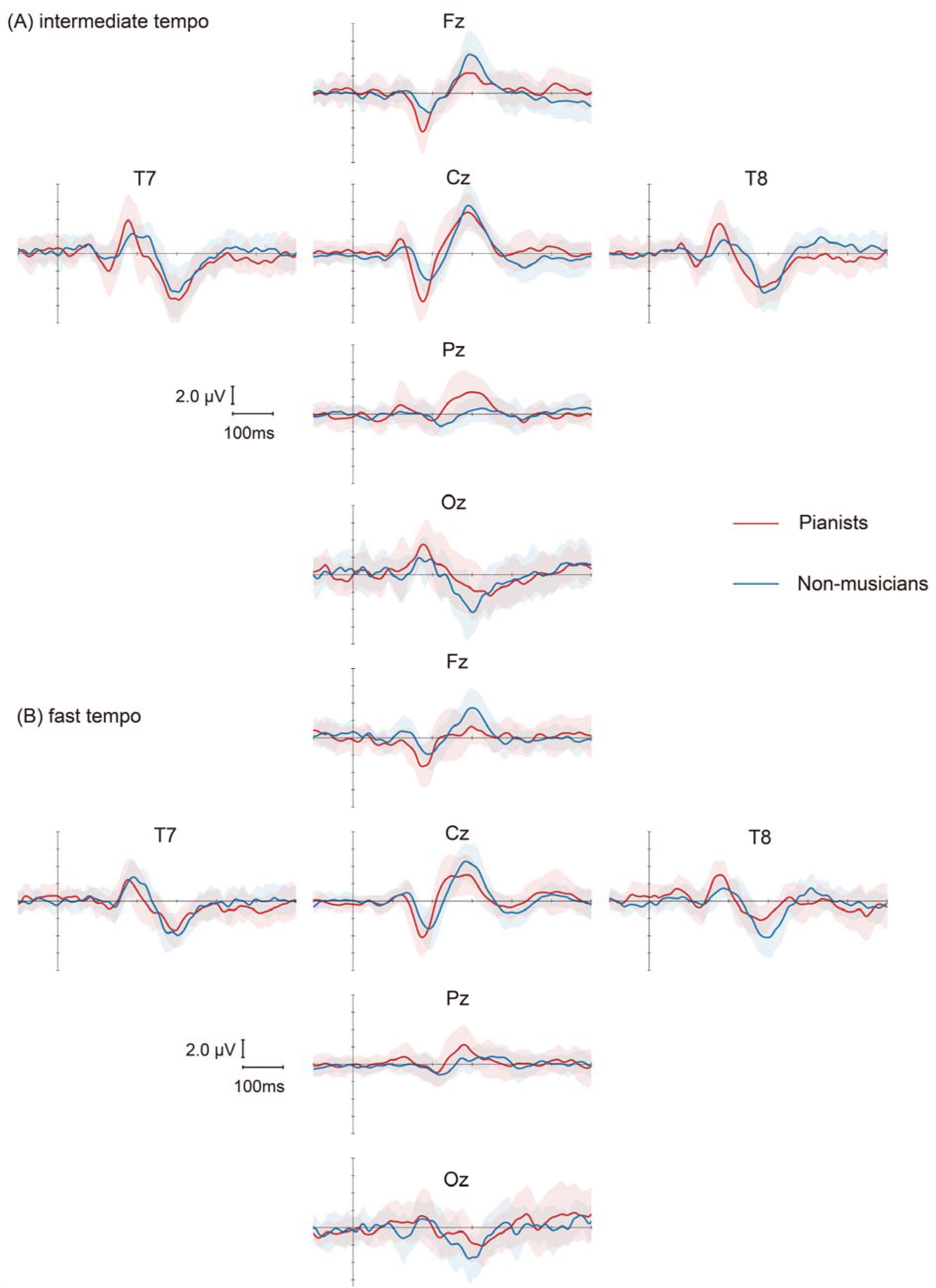
The changes in grand-averaged ERPs (perturbed - unperturbed conditions) over keystrokes after the tone production was transiently delayed. The red and blue lines represent the pianists and non-musicians, respectively. The shaded area around the line represents one standard deviation across the participants. (A) and (B) represent a comparison between the pianists and non-musicians when playing at the intermediate and fast tempo, respectively. Time zero was defined as the moment when the depression of the key with delayed auditory feedback was initiated. The negative component N180 was elicited at around 180 ms following the tone delay (= a vertical line), whereas the positive component P300 was elicited at around 300 ms in both groups.

For the amplitude of N180, the three-way ANOVA with group, tempo, and channel showed a significant interaction effect between group and tempo (Table 2). Post hoc tests, however, revealed no significant differences between groups and between tempi.

**Table 2.**
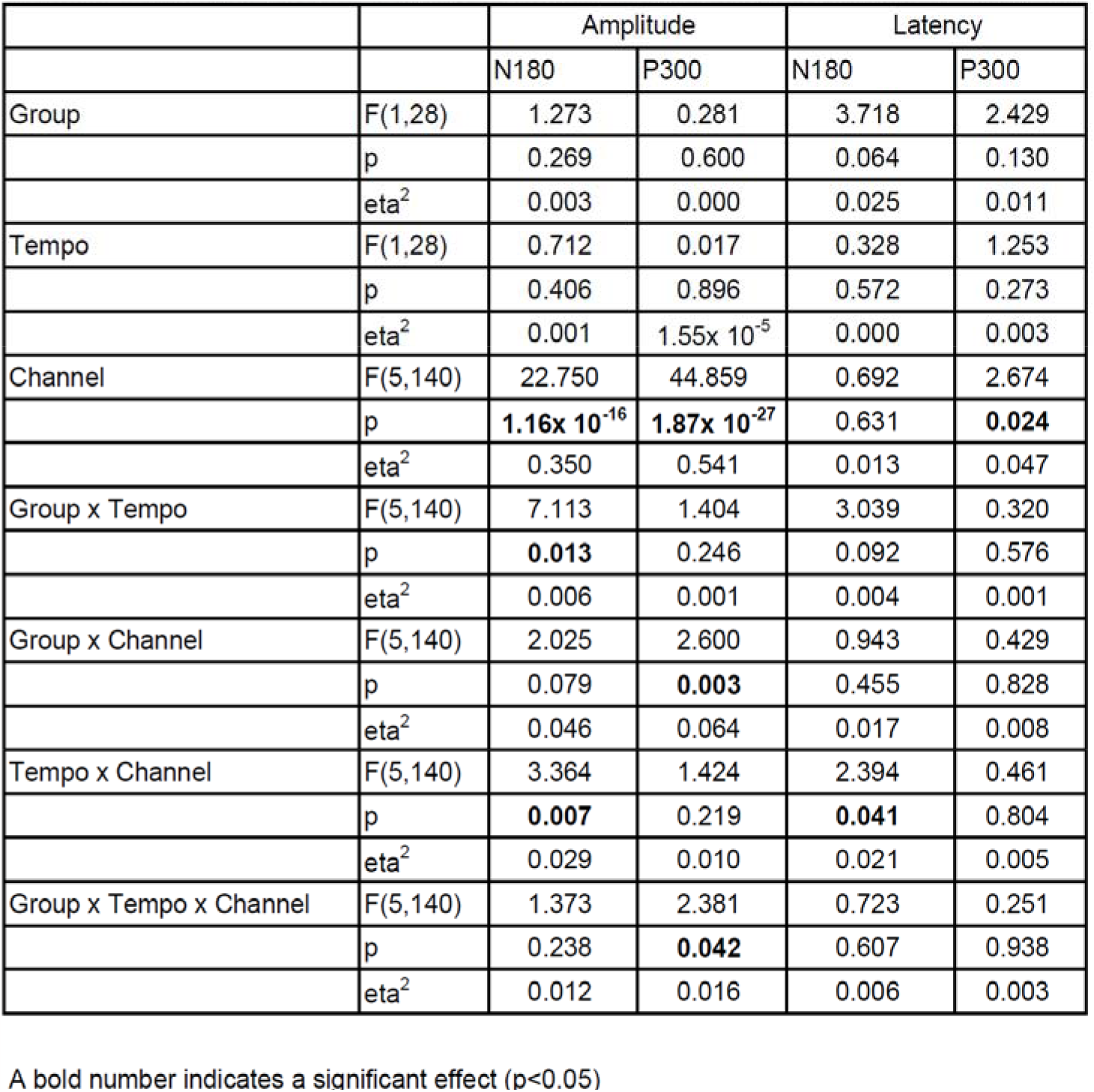
Summary of the three-way repeated mixed-design ANOVA with factors of Group, Tempo and EEG channels.

|  |  | Amplitude |  | Latency |  |
| --- | --- | --- | --- | --- | --- |
|  |  | N180 | P300 | N180 | P300 |
| Group | F(1,28) | 1.273 | 0.281 | 3.718 | 2.429 |
|  | p | 0.269 | 0.600 | 0.064 | 0.130 |
|  | eta<sup>2</sup> | 0.003 | 0.000 | 0.025 | 0.011 |
| Tempo | F(1,28) | 0.712 | 0.017 | 0.328 | 1.253 |
|  | p | 0.406 | 0.896 | 0.572 | 0.273 |
|  | eta<sup>2</sup> | 0.001 | 1.55x 10<sup>-5</sup> | 0.000 | 0.003 |
| Channel | F(5,140) | 22.750 | 44.859 | 0.692 | 2.674 |
|  | p | **1.16x 10<sup>-16</sup>** | **1.87x 10<sup>-27</sup>** | 0.631 | **0.024** |
|  | eta<sup>2</sup> | 0.350 | 0.541 | 0.013 | 0.047 |
| Group x Tempo | F(5,140) | 7.113 | 1.404 | 3.039 | 0.320 |
|  | p | **0.013** | 0.246 | 0.092 | 0.576 |
|  | eta<sup>2</sup> | 0.006 | 0.001 | 0.004 | 0.001 |
| Group x Channel | F(5,140) | 2.025 | 2.600 | 0.943 | 0.429 |
|  | p | 0.079 | **0.003** | 0.455 | 0.828 |
|  | eta<sup>2</sup> | 0.046 | 0.064 | 0.017 | 0.008 |
| Tempo x Channel | F(5,140) | 3.364 | 1.424 | 2.394 | 0.461 |
|  | p | **0.007** | 0.219 | **0.041** | 0.804 |
|  | eta<sup>2</sup> | 0.029 | 0.010 | 0.021 | 0.005 |
| Group x Tempo x Channel | F(5,140) | 1.373 | 2.381 | 0.723 | 0.251 |
|  | p | 0.238 | **0.042** | 0.607 | 0.938 |
|  | eta<sup>2</sup> | 0.012 | 0.016 | 0.006 | 0.003 |

A bold number indicates a significant effect (p<0.05)

The amplitude of P300 results showed a significant three-way interaction effect (Table 2). Post hoc tests revealed significant differences between the groups at the intermediate tempo at the Pz, Oz, and Fz channels, showing a larger P300 amplitude in the pianists. There was also a significant difference between the tempi at the T8 channel of the pianists.

For the latency of ERPs, neither interaction effects nor main effects were significant at both N180 and P300 (Table 2).

### Multiple-regression analysis

Table 3 summarizes the results of the Ridge regression analyses, which explained the changes in each of the inter-keystroke intervals and the key descending velocity when playing at the intermediate tempo according to ERPs in response to the delayed tone production. The values denoted in Table 3 indicate partial regression coefficients. Figure 4 represents the coefficients derived from the regression analysis, plotted on a topographic map related to the behavioral responses to the perturbation.

**Fig. 4.**
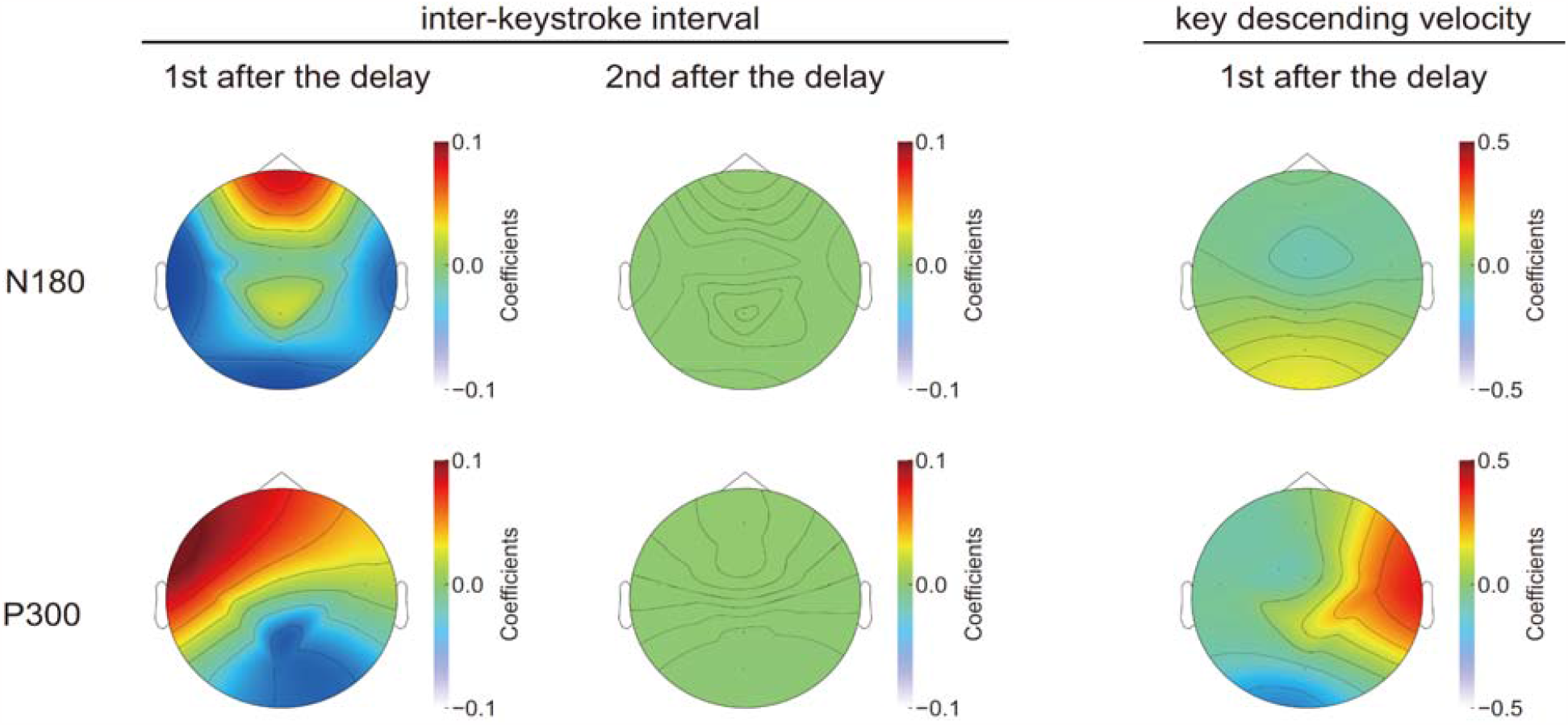
Topographic distribution map representing the coefficients derived from the regression analysis. The distribution is displayed from the top, looking down at the head. The distribution was visualized to intuitively grasp the results outlined in Table 3.

**Table 3.** Summary of the Ridge regression analyses between the individual behavioral features of the keystrokes and ERPs at the intermediate tempo.

|  |  | Interval |  | Velocity |
| --- | --- | --- | --- | --- |
|  |  | 1st after | 2nd after | 1st after |
| N180 | Fz | 0.064 | 0.001 | -0.053 |
|  | Cz | -0.008 | -0.001 | -0.104 |
|  | Oz | -0.039 | 0.000 | 0.109 |
|  | T7 | -0.056 | -0.001 | -0.037 |
|  | T8 | -0.040 | 0.000 | -0.054 |
|  | Pz | 0.020 | 0.001 | 0.010 |
| P300 | Fz | 0.073 | 0.001 | -0.024 |
|  | Cz | 0.027 | 0.001 | -0.053 |
|  | Oz | -0.051 | -0.001 | -0.115 |
|  | T7 | 0.084 | 0.000 | -0.068 |
|  | T8 | 0.012 | 0.000 | 0.341 |
|  | Pz | -0.052 | -0.001 | 0.102 |
| $R^2$ | | 0.209 | 0.003 | 0.385 |

Interval: inter-keystroke interval
Velocity: key descending velocity
A sign indicates whether each dependent variable covaried positively or negatively with the independent variable.

Coefficients of the first inter-keystroke interval following the perturbation revealed that the prolonged interval, which represents the disruption of the local tempo, was associated with the decrease of both the amplitude of N180 in the central, temporal, and occipital regions and the amplitude of P300 in the left temporal and parietal regions, and with the increase in the amplitude of P300 in the frontal and occipital regions.

With respect to the first key descending velocity after the perturbation, the coefficients revealed that the stronger strike at the first keystroke after the tone delay was related to larger P300 elicited from the right temporal region. This result indicates an association of the right temporal region with movement adaptability to the perturbation.

For the second inter-keystroke interval after the perturbation, both the coefficient and R^2^ were close to zero, indicating no relationship between the behavioral responses at this event and the ERPs.

## Discussion

In this study, we aimed to address differences in the effects of auditory perturbation on motor and electrophysiological responses across individuals with different skill levels by leveraging the novel behavioral and neurophysiological measurement system during fast piano performance. Behaviorally, non-musicians exhibited prolonged inter-keystroke intervals after auditory perturbation. Conversely, expert pianists showed minimal changes in the inter-keystroke intervals after the perturbation, which indicates the robustness of the performance. They even appeared to compensate promptly for the tone delay by speeding up the local tempo. Furthermore, pianists were capable of adjusting the key descending velocity immediately after the tone delay, probably as a way of elevating sensory gain in motion. The observation indicates that expert pianists possess a superior ability to adapt their actions online in response to auditory perturbation. Neurophysiologically, ERPs in the N180 and P300 components from the onset of the keystroke were elicited in both groups following the delayed tone production. Especially, P300 (i.e., around the parietal region) associated with aspects of cognition and efficient information processing in the brain was more sensitive to adaptability of skillful actions. Specifically, expert musicians exhibited a significantly larger response in the Pz electrode compared to the non-musicians. The central question of the present study was to ascertain whether there exists a neural correlate explaining the high adaptability of the expert musicians to the perturbation. Our regression model revealed a close association of ERPs at both N180 and P300, specifically in the frontal and temporal regions, with the alterations in the inter-keystroke interval and key descending velocity following the perturbation, respectively. Taken together, for the first time, we provide novel evidence suggesting that neural activities related to cognition and auditory processing play a vital role in adaptability and robustness of skillful behavior in expert musicians during fast sequential actions.

### Expertise-dependent robustness and adaptability of skillful behavior

Expert musicians possess the capacity to dynamically and flexibly adjust their keystrokes with precision in terms of both force and tempo. This exceptional ability results not only from their finely-tuned sensorimotor control but also from their cognitive capabilities, which include the capacity to discern subtle sensory perturbations. This cognitive skill enables them to execute skillful motor actions seamlessly during piano performances. However, this proves to be fairly challenging for non-musicians. It is not an exaggeration to assert that this difference in adaptability leads to variations in skillful performance specifically of expert musicians. While variations in the degree of adaptability exist, the fundamental ability to detect disruptions and errors plays a pivotal role in motor adaptation (Tan et al., 2014; Seidler et al., 2015; Uehara et al., 2018, 2019). Through prolonged and intensive training leading to the acquisition of advanced musical skills, expert pianists may have honed their capacity for sensory-related error detection and the ability to promptly correct ongoing actions. This assumption is corroborated by our behavioral observations, which demonstrate that expert pianists exhibited minimal changes in inter-keystroke intervals immediately after auditory perturbations and employed a strategy to promptly recover from the tone delay (see Figure 2A). Consequently, they can play robustly against the perturbation, unlike non-musicians. The findings corroborate with previous behavioral studies reporting that expert pianists possess a higher level of robustness against temporal perturbations (van der Steen et al., 2014) and the ability to compensate for the delay by advancing the subsequent keystroke after the delay to maintain the overall tempo of the musical performance (Furuya and Soechting, 2010).

Interestingly, in response to the delay, non-musicians pressed the key more softly, whereas pianists pressed the key harder (see Figure 2B). This result suggests that pianists proactively attempted to correct the delayed sensory consequence immediately. One plausible explanation is that expert pianists may elevate sensory gain to rely more on sensory-feedback control that allows for exploring the optimal action and/or maintaining online control of movements immediately after being perturbed. This is because an internal model in the nervous system, which predicts sensory consequences of actions, was disrupted by unexpected auditory feedback, which then increases reliance on sensory feedback control according to the framework of the optimal feedback control principle (Todorov, 2004; Scott, 2016). Following the auditory perturbation, pianists struck the key stronger, which may propose two putative mechanisms. They may attempt to receive stronger proprioceptive feedback from the finger. Alternatively, a stronger keystroke elicits a loud sound that can be utilized to correct ongoing actions through augmenting auditory feedback gain in motion. Consequently, we presume that pianists excel in terms of robustness against and adaptability to perturbation through undergoing intensive musical training.

### Functional significance of ERPs and their impact on skillful behavior

In this study, we focused on EEG-based ERPs as a candidate neural signature that emerges in response to transient sensory perturbation (Maidhof et al., 2010; Strübing et al., 2012; Sammler et al., 2013; Bianco et al., 2016). ERPs are characterized by specific patterns of EEG activity in response to particular stimuli or events, encoding valuable neural information about the processing of cognitive, sensory, and motor functions (Polich, 2007; van Dinteren et al., 2014; Hirano et al., 2020; Iwane et al., 2023). This neural index enables us to shed light on cognitive and sensorimotor processes. ERPs can be analyzed to identify components based on their latency from the onset of particular stimuli or events (Squires et al., 1975; van Dinteren et al., 2014; Iwane et al., 2023). Among them, we drew attention to the observations of the N180 and P300 components. The negative potential around 180 ms has been suggested to be the error-related negativity (ERN), which is an index of the discrepancy between expected and actual occurrence of events (Maidhof et al. 2010; Proverbio et al. 2017) as well as performance monitoring (Iwane et al., 2023). ERN often emerges from the anterior cingulate cortex (ACC) located in the frontal cortex when individuals receive an erroneous action (Ullsperger and von Cramon, 2001; Brázdil et al., 2002). A human lesion study also demonstrated the impaired ability of error monitoring due to lesions in the ACC (Maier et al., 2015). Thus, the ACC has been a candidate brain region responsible for error detection. Recent human EEG studies have provided further evidence that ERNs code not only the awareness of errors but also the magnitude of errors in their amplitude (Spinelli et al., 2018; Iwane et al., 2023). As for the P300 component, this long-latency response is one of the most commonly reported components. Empirical evidence indicates that the P300 component is associated with metacognition including error monitoring and confidence (Boldt and Yeung, 2015). In contrast to the ERN, P300 is more closely associated with subjective error awareness, i.e., the subjective judgment of response accuracy (Steinhauser and Yeung, 2010). One of the important comparisons in our dataset is between-group differences, reflecting expertise-dependent proficiency. We found that pianists exhibited a significantly larger P300 response at the Pz electrode underneath the mid-central brain region compared to non-musicians, but not for the N180 response. This indicates that pianists have a superior ability to subjective error awareness in response to perturbation than non-musicians. Moreover, given that the P300 response is associated with error-related metacognition, expert musicians may be superior in terms of instantaneous decision-making to correct sensorimotor disruption.

Based on the aforementioned neurophysiological group comparisons, we presume that the P300 response is better characterized as reflecting expertise-dependent robustness and adaptability of skillful behavior. To address this, ridge regression analyses were employed in this study. This approach unmasked the neural correlate to the alterations of inter-keystroke interval and key descending velocity after auditory perturbation, and these neural correlations emerged across the extensive cortical regions rather than localized brain regions (see Figure 4). Additionally, the inter-keystroke interval at the first strike after (1st after) the perturbation was explained by both ERPs, unlike the 2nd strike after it (R^2^ was 0.003). This result indicates that neural activities related to error correction emerge within less than 500ms and vanish quickly. Our high-speed behavioral and neurophysiological measurement system allowed for successfully capturing this unique phenomenon.

The specific behavioral trait observed only in pianists that should be discussed here together with our novel results of ERPs includes the sensory gain adjustment as mentioned above. We argued that proprioceptive, auditory, or both may aid in adapting motor actions to auditory perturbation. Thus, a hierarchical structure of the multi-sensory system regarding the adaptability and robustness of sensorimotor control in response to perturbation may exist. Our regression analysis provides a clue in favor of this assumption. We found that key descending velocity co-varied positively with P300 emerging from the right temporal region (Figure 4 right panel), but not the left somatosensory region (contralateral to the task side). If proprioceptive feedback is more involved following the perturbation, the left sensory region may also covary functionally with the descending velocity. We therefore suggest that the emphasis on the auditory feedback stemming from the increased key descending velocity plays a more dominant role in gain control than proprioceptive feedback, which has been unknown. We might extend this interpretation to understanding hierarchical representations of sensation, indicating that auditory-motor integration may reside in a higher hierarchy than somatosensory-motor integration in expert pianists. To test this novel hypothesis requires further studies.

### Study limitation

The present study has a limitation. We designed our experiment to reveal the neural correlates of expertise-dependent robustness and adaptability of skillful behavior. Therefore, it is still debatable whether a causal relationship between neural activities and behavioral characteristics exists or not. This limitation must be considered when interpreting our results. Our next step will be to design a causal approach using a virtual lesion approach with non-invasive brain stimulation or EEG-based neurofeedback. However, given that the adaptation of skillful performance is completed instantaneously, it is very challenging to manipulate target brain activities under high-speed actions.

## Acknowledgement

This work was supported by JST, Moonshot R&D Grant Number JPMJMS2012.

